# Stage-Specific MCM Protein Expression in *Trypanosoma cruzi*: Insights Into Metacyclogenesis and G1 Arrested epimastigotes

**DOI:** 10.1101/2025.03.05.641675

**Authors:** Bruno Alves Santarossa, Évelin Mariani, Artur da Paixão Corrêa, Fernanda C. Costa, Martin C. Taylor, John M. Kelly, Maria Carolina Elias, Simone Guedes Calderano

**Affiliations:** Cell Cycle Laboratory, Butantan Institute, Av. Vital Brazil 1500, São Paulo, Brazil; Department of Infection Biology, London School of Hygiene and Tropical Medicine. Keppel Street, London WC1E 7HT, UK

**Keywords:** mini-chromosome maintenance, MCM, *Trypanosoma cruzi*, metacyclogenesis, replication control, G0, G1 arrest, cell cycle arrest

## Abstract

*Trypanosoma cruzi* is a protozoan parasite that is the etiological agent of Chagas disease, which is endemic to Latin America with reported cases in non-endemic regions such as Europe, Asia, and Oceania due to migration. During its lifecycle, *T. cruzi* alternates between replicative and non-replicative infective lifeforms. Metacyclogenesis is the most studied transition in which replicative epimastigotes differentiate into infective metacyclic trypomastigotes inside the gut of the triatomine vector. This early-branching organism presents a divergent pre-replication complex (pre-RC) where the only conserved component is the MCM2-7 protein family. Given the role of pre-RC components in cell cycle regulation, we investigated whether MCM expression and location could be involved in proliferation control in epimastigotes and during metacyclogenesis. Our findings reveal that MCMs are consistently expressed and localized to the nucleus throughout the epimastigote cell cycle, including in G1/G0-arrested cells. However, MCM proteins are degraded during metacyclogenesis as cells enter the G0 state, marking the transition to replication arrest. Therefore, epimastigotes arrested in G1/G0 can either maintain MCM expression and resume the cell cycle when conditions become favorable, or they can undergo metacyclogenesis, exiting the cell cycle and entering a G0 state, where MCMs are degraded as part of the replication repression mechanism.

## Introduction

Cellular commitment to the cell cycle is strongly influenced by environmental conditions, such as nutrient availability. Under favorable conditions, cells pass through the G1 checkpoint, and progress irreversibly through the cell cycle, ultimately generating two daughter cells (Blagosklonny and Pardee, 2002, Johnson and Skotheim, 2013). Accurate DNA replication is essential during this process. DNA replication is initiated at specific chromosomal sites named origins of replication. During the cell cycle, two main events must occur at these origins: licensing and firing. The licensing process occurs in late mitosis and early G1 phases and is accomplished by assembling the pre-Replicative Complex (pre-RC). The Origin Recognition Complex (ORC1-6) recognizes and binds to the origins of replication and allows the subsequent loading of Cdt1 and Cdc6, which carries the Mini-Chromosome Maintenance complex (MCM2-7), completing pre-RC assembly (Stoeber et al., 2001). The MCM2-7 complex has a ring-shaped structure with helicase activity that is only activated after CDC45 and GINS recruitment, at the beginning of the S phase, when origins of replication are fired (Im et al., 2009). To guarantee that each origin of replication is fired only once per cell cycle, the pre-RC components are subject to degradation, detachment from chromatin, and export from the nucleus, thus avoiding DNA re-replication (Arias and Walter, 2007). In both budding yeast and fission yeast, all ORC subunits remain bound to chromatin throughout the cell cycle, however, Orc2 and Orc6 are phosphorylated and lose their capacity to load the MCM2-7 complex (Fujita et al., 1998, Diffley et al., 1995, Liang and Stillman, 1997, Lygerou and Nurse, 1999). In metazoan, pre-RC subunits are modified in a cell cycle-specific manner that controls chromatin affinity, cellular localization and/or stability (DePamphilis, 2005). For instance, Orc1 can be ubiquitinated and degraded during the S phase in HeLa cells (Méndez et al., 2002), or detached from chromatin and accumulated in the cytoplasm of CHO cells (Saha et al., 2006). Cdc6 is phosphorylated and degraded during the S phase in yeast (Drury et al., 2000, Jallepalli et al., 1997) and during mitosis in metazoan, but it also can be exported to cytoplasm in metazoans (Petersen et al., 2000, Kim and Kipreos, 2008). Cdt1 is exported from the nucleus in budding yeast during the S phase (Tanaka and Diffley, 2002) and is degraded in fission yeast and metazoan, also in S phase (Blow and Dutta, 2005, Machida et al., 2005). Additionally, in metazoan, Cdt1 can be inhibited by the binding protein geminin (Kim and Kipreos, 2007). In budding yeast, MCM2-7 is exported from the nucleus, and in fission yeast and metazoan, MCM2-7 remains nuclear but is detached from chromatin at the end of DNA replication. Therefore, regulation of the pre-RC complex is conserved among eukaryotes, with distinct mechanisms observed across different systems.

When cells face unfavorable conditions, such as nutritional deprivation, the restriction point is not crossed and the cells exit the cell cycle (G0 phase). At this point, cells can either remain quiescent until the cell cycle resumes under favorable environmental conditions or they can terminally differentiate, entering a permanent non-proliferative state where the cell cycle does not resume (Sagot and Laporte, 2019), (Matson and Cook, 2017, Fiore et al., 2018). In both situations, the pre-RC are downregulated (Blow and Hodgson, 2002, Stoeber et al., 2001), but in terminally differentiated cells some pre-RC can be completely degraded (Carroll et al., 2018).

*Trypanosoma cruzi* is a protozoan parasite that causes Chagas’ disease, a potentially life-threatening illness endemic to Latin America with an estimated 6 to 7 million people infected (WHO, 2024). During its lifecycle, *T. cruzi* transitions from replicative to non-replicative lifeforms inside the mammalian host (amastigote and bloodstream trypomastigotes, respectively), and the triatomine insect vector (epimastigote and metacyclic trypomastigotes, respectively)(Martín-Escolano et al., 2022). In the vector, the process is called metacyclogenesis, and nutritional stress plays a crucial role in triggering the transition to infectious trypomastigotes (Melo et al., 2020). *T. cruzi*, *Trypanosoma brucei* and *Leishmania spp.* are the three main human-infectious trypanosomatids, and their pre-replication complexes differ from other eukaryotes. The ORC is divergent (Godoy et al., 2009, Tiengwe et al., 2012, Marques et al., 2016), no Cdt1 and Cdc6 equivalents have been found, but all components of the MCM2-7 complex are conserved (Dang and Li, 2011, Tiengwe et al., 2012, da Silva et al., 2017). Since MCM2-7 appeared to be the only pre-RC conserved component, we questioned whether MCM expression and location could be involved in proliferation control in *T. cruzi*. For this, we analyzed replicating and stationary epimastigotes, and those undergoing metacyclogenesis. We have previoulsy found that subunit 7 of the MCM2-7 complex is only expressed in replicative lifeforms and is abolished in non-replicative trypomastigotes (metacyclic and bloodstream) (Calderano et al., 2014). Here, we investigated other MCMs subunits and found that MCMs are expressed and localized inside the nucleus of replicating and stationary epimastigotes, and that degradation only occurs during metacyclogenesis, when cells are arrested in G0. While epimastigotes retain an ability to re-enter the cell cycle, MCMs remain expressed, and cytoplasmic export is not involved as a regulatory mechanism. Degradation occurs only after differentiation into non-proliferative metacyclic trypomastigotes, and MCM expression is restored after differentiation to the replicative lifeform amastigotes within infected host cells.

## Methodology

### Cell Culture

#### Epimastigotes

*T. cruzi* strains CL Brener and Dm28c, were maintained as epimastigotes in LIT medium (Camargo, 1964) supplemented with 10% fetal bovine serum (FBS), 60 µg/mL of penicillin,100 µg/mL of streptomycin, and incubated at 28°C.

#### Hydroxyurea synchronization

Epimastigote cultures were synchronized with Hydroxyurea (HU) as described previously (Galanti et al., 1994). Briefly, epimastigote cultures were diluted in fresh medium to a final concentration of 3×10^6^/mL. After 24h, HU was added at a final concentration of 20 mM and incubated for 24h at 28 °C. Cells were washed three times with PBS and suspended in a fresh medium. Aliquots were collected at 0h, 4h, 6h, and 8h after HU removal, which were fixed for FACs analysis and total protein extraction.

#### Metacyclogenesis

The *in vitro* metacyclogenesis process was based on Contreras et al., 1985, with some changes. Epimastigote cultures in the stationary growth phase (∼10^8^ parasites/mL) were stressed in TAU medium (190 mM NaCl, 8 mM phosphate buffer pH 6.0, 17 mM KCl, 2 mM CaCl2, 2 mM MgCl2) at 5×10^8^ parasites/mL for 2h at 28 °C. Afterwards, they were diluted to 5×10^6^ parasites/mL in TAU3aaG (TAU supplemented with 10 mM L-proline, 50 mM L-sodium glutamate, 2 mM L-sodium aspartate and 10 mM D-glucose) and incubated for 5 days at 28 °C, in 5% CO2. To determine the percentage of differentiation, cells were fixed on a slide and stained with DAPI. Nucleus, kinetoplast, and cellular morphology were analyzed to determine epimastigotes, intermediates and fully differentiated trypomastigotes (Gonçalves et al., 2018).

#### *In vitro* infective cycle of *T. cruzi*

A total of 2.5 x 10^5^ LLC-MK2 cells were seeded in a 175 cm^2^ flask with 40 mL of DMEM supplemented with 10% FBS and incubated at 37°C in 5% CO2. After 24h, 10^6^ TCTs (Tissue Cultured Trypomastigotes) were added to the LLC-MK2 culture and incubated for 24h under the same conditions. The remaining TCTs were removed by washing with PBS and fresh medium added. After 5-6 days, TCTs were collected from the medium, following their egress from host cells.

### CRISPR/Cas9

Gene editing was performed as described (Costa et al., 2018) targeted at the following genes in the CL Brener (MCM2: TcCLB.506933.40, MCM3: TcCLB.511109.100, MCM4: TcCLB.511127.140, MCM5: TcCLB.508647.140, MCM6: TcCLB.507527.30) and Dm28c strains (MCM6: C4B63_6g251, MCM7: C4B63_80g19). Briefly epimastigotes expressing the Cas9 enzyme and T7 RNA polymerase were transfected using program X-014 from Nucleofactor2b (Lonza) (Burkard et al., 2007) and the electroporation buffer (90 mM sodium phosphate, 5 mM potassium chloride, 0.15 mM calcium chloride, 50 mM Hepes, pH 7.2).

CRISPR/Cas9 transfections were carried out with two PCR products (∼5 µg of each) representing single guide RNA (sgRNA) and donor DNA as described previously (Costa et al., 2018). Donor DNA to insert 3 copies of mNeonGreen, 6 copies of Myc and blasticidin resistance gene were amplified from pPOTv6-blast-3Myc::3mNG::3Myc plasmid (Paterou et al., 2023). Donor DNA to insert 3 copies of Myc and puromycin resistance gene were amplified from pMOTag (Lander et al., 2017).

**Table 1:**
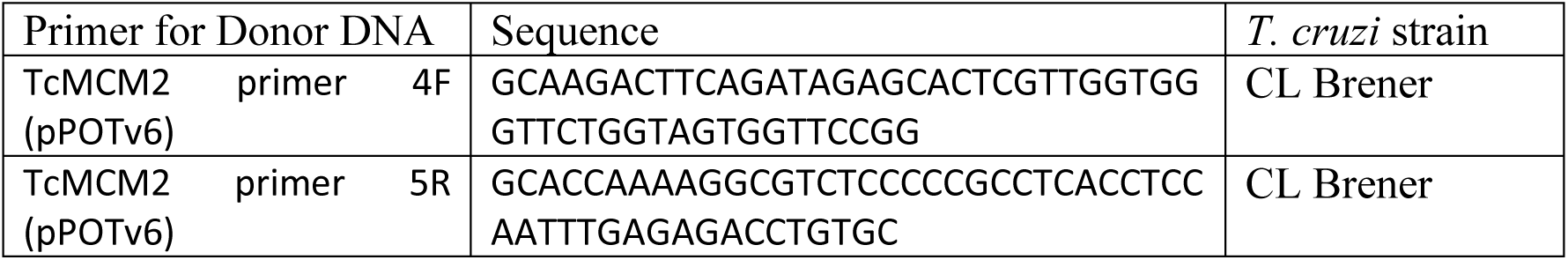
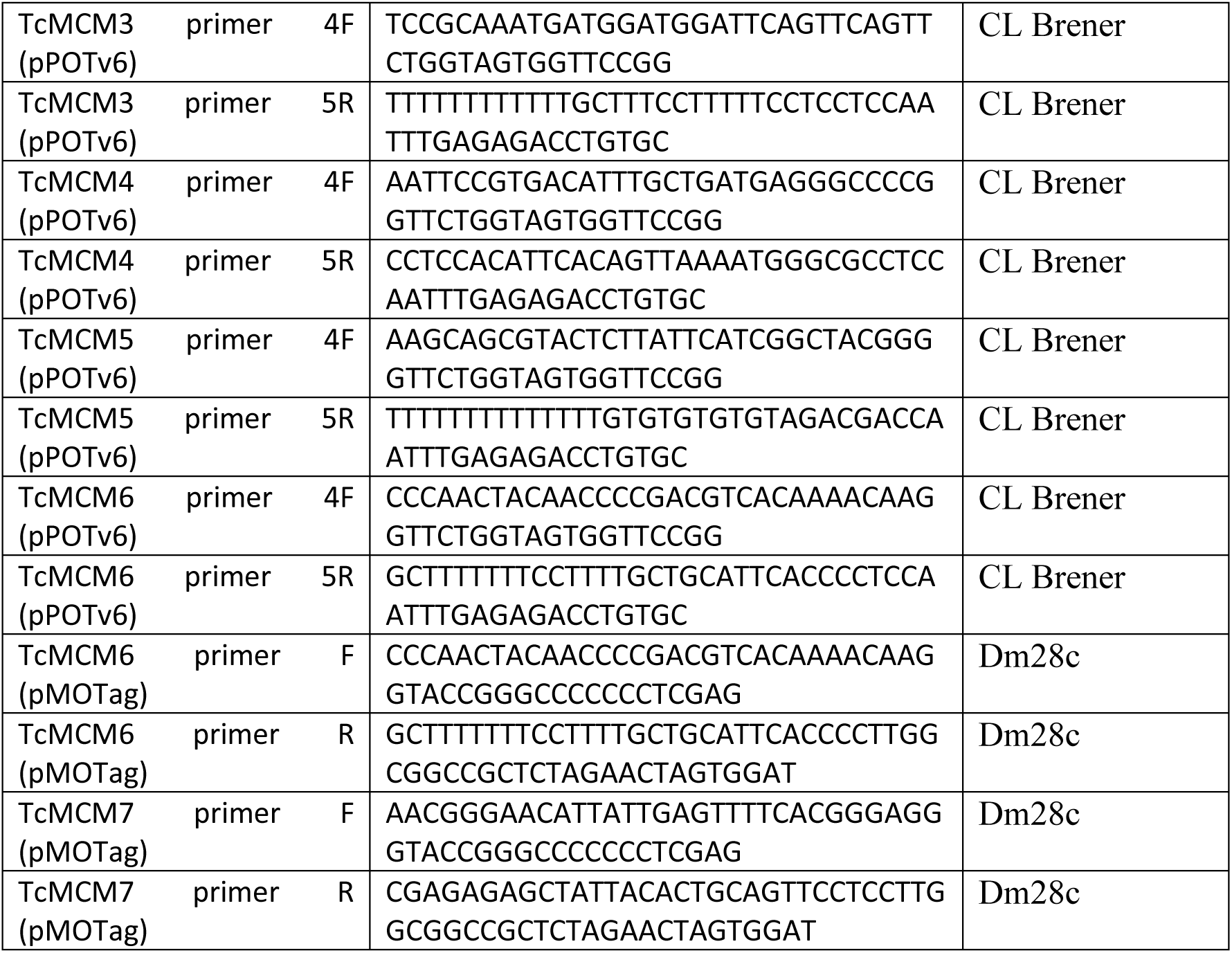
List of primers used to amplify donor DNA.

**Table 2:**
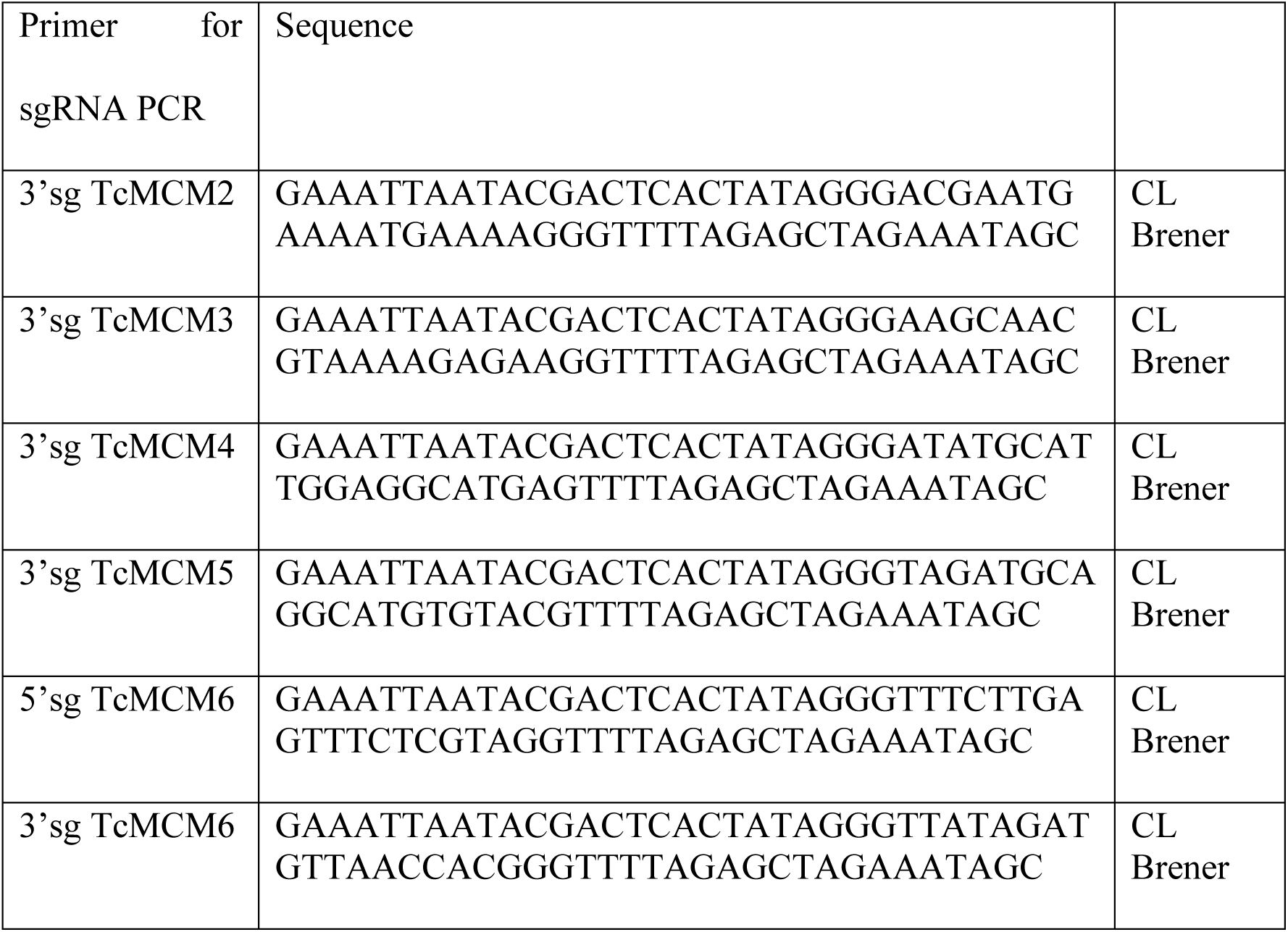
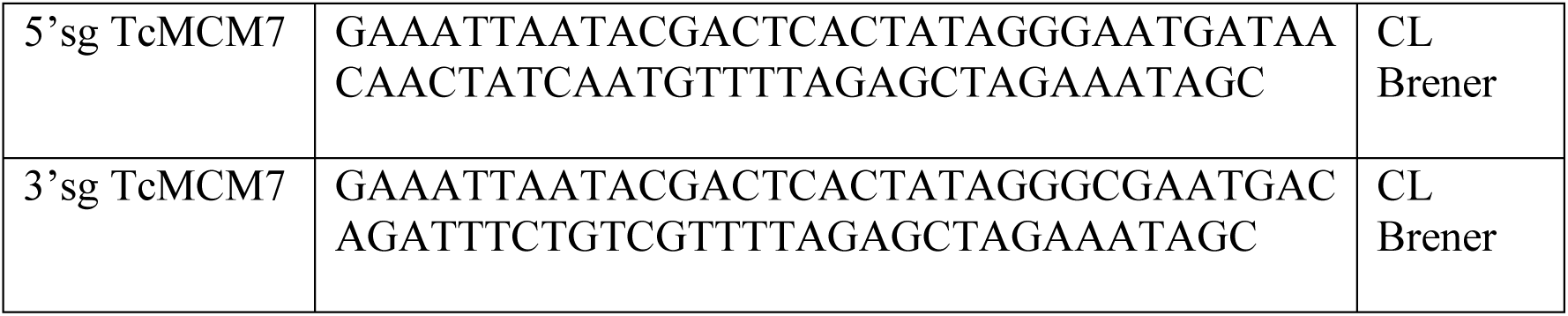
List of primers used to obtain PCR product for sgRNA.

### Epimastigotes cloning

Epimastigotes were counted in a Neubauer chamber and diluted to a final concentration of 1 parasite/mL of LIT medium. Then 200 µL aliquots of this diluted culture were applied to each well of a 96 well plate and incubated at 28 °C.

### Immunofluorescence

#### Extracellular life-cycle stages

For immunofluorescence assays, 3×10^6^ cells were applied to poly-D-lysine-coated glass slides, incubated at room temperature for 5 minutes and washed with PBS. Then cells were fixed with 2% paraformaldehyde for 10 minutes, washed twice with PBS and permeabilized with PBS/0.3% Triton X-100 for 5 minutes. After washing twice with PBS, the cells were blocked for 1h in PBS containing 3% of bovine serum albumin, 1% bovine gelatin and 50% FBS. The cells were then washed twice with PBS and incubated at room temperature for 4h in PBS/3% BSA with mouse anti-Myc (1:1000) (Myc-Tag 9B11 Mouse mAB, Cell Signalling). The cells were then washed 5x with PBS and incubated for 1h at room temperature with PBS/BSA containing anti-mouse 488 plus (1:1000) (Thermo Fisher Scientific). After several washes with PBS, the slide was mounted using Vecta-Shield with DAPI (VectorLabs). Images were acquired using a 100× 1.35NA lens and cell F software in an Olympus BX51 microscope (Tokyo, Japan). Brightness and contrast were adjusted using Photoshop. Raw images of control and edited cell lines are presented in Supplementary Figure 2.

#### EdU staining

Cells were incubated for 30 minutes with 5-ethynyl-2′-deoxyuridine (EdU) at a final concentration of 100 µM, washed 3x with PBS, and processed as above for microscopy, as previously described. EdU detection followed the manufacturer’s instructions (Click-iT™ EdU Cell Proliferation Kit for Imaging, Alexa Fluor™ 647 dye-Thermo Fisher Scientific). Images were acquired through a z-series of 0.2 μm using a 100× 1.35NA lens and cell R software in an Olympus IX81 microscope. Deconvolution was performed using AutoQuant X software.

#### Intracellular Amastigotes

LLC-MK2 cells (5 × 10³) were seeded on 13 mm round coverslips in a 12-well plate using DMEM supplemented with 10% FBS. The cells were incubated for 24h at 37 °C with 5% CO₂. Cells were infected with Dm28c strain trypomastigotes at an MOI of 1:10. After 24h, the medium was removed, and the cells washed with PBS to eliminate remaining parasites. Fresh DMEM with 10% FBS was then added to the wells. Following a 48h incubation period, the cells underwent immunofluorescence staining. The steps for fixation, permeabilization, blocking, and antibody incubation were performed as previously described.

### Western Blotting

RIPA Lysis and Extraction Buffer (Thermo Fisher Scientific) supplemented with a protease inhibitor cocktail (Pierce^TM^ Protease Inhibitor Tablets, EDTA free-Thermo Fisher Scientific) and a phosphatase inhibitor cocktail (Halt^TM^ Phosphatase Inhibitor Cocktail-Thermo Ficher Scientific) were used to extract total proteins, using 20 µL of RIPA buffer for each 10^7^ cells. Total protein extract concentration was determined using a Pierce^TM^ BCA Protein assay kit (Thermo Fisher Scientific).

### Western blotting band quantification using Photoshop

Western blotting images were acquired using a UVITEC system (Cambridge), employing automatic exposure times to prevent band saturation. Subsequently, images were analyzed using Photoshop software to quantify band intensity. Signal (anti-Myc) and control (anti-flag) images were juxtaposed in the same JPEG file, enabling uniform band quantification under consistent parameters. To ensure accurate measurement, the rectangle selection tool was utilized to delineate the band area. A consistent selection size was applied across all bands under analysis, encompassing both the anti-Myc signal and control bands. Additionally, a nearby region of each band was selected as the background signal. The intensity of each selected area (anti-Myc, anti-Flag, and background) was quantified to facilitate the calculation of the relative expression levels of MCM proteins. The background intensity was subtracted from the intensities of the anti-Myc and anti-Flag signals to correct for non-specific signal contributions. The corrected anti-Myc intensity was normalized to the corrected anti-Flag intensity by dividing the anti-Myc intensity by the anti-Flag intensity. This step accounts for any variability in the total protein load or other experimental inconsistencies. The relative expression levels were determined by dividing the normalized anti-Myc/anti-Flag ratio for each sample by the normalized anti-Myc/anti-Flag ratio of a reference sample (e.g., a control or baseline sample). This provided a relative measure of MCM expression across different samples. Data visualization and statistical analyses were performed using GraphPad Prism software. Graphs were created to illustrate the results, and appropriate statistical tests were applied to assess the significance of the differences observed.

### Flow Cytometry

#### Propidium iodide

10^7^ cells were pelleted, washed twice with PBS and fixed by resuspending in 1 mL of cold 70% ethanol/30% PBS and kept at –20 °C for at least 4h. Fixed cells were pelleted, washed twice with PBS and resuspended in 500 µL PBS containing RNAse A (Invitrogen) and propidium iodide (100 µM final concentration) (Thermo Fisher Scientific). The cells were incubated at 37 °C for 30 minutes and analyzed by using Attune NxT (Life Technologies) flow cytometer.

#### EdU incorporation

after 30 minutes incubation with 100 µM EdU (final concentration) (Thermo Fisher Scientific), 10^7^ cells were pelleted, washed 3x with 1 mL of PBS. The cell pellet was then fixed and EdU detection performed as previously described and analyzed using Attune NxT flow cytometer.

## Results

### MCM6-Myc and MCM7-Myc expression is constitutive throughout epimastigote cell cycle

We utilized the CRISPR/Cas9 system to tag MCM proteins in *T. cruzi* CL Brener and DM28c epimastigotes. Specifically, we inserted three copies of mNeonGreen and six copies of a c-Myc epitope at the 3’ end of the MCM 2, 3, 4 and 6 genes in the CL Brener strain (Figure 1 A), and three copies of the c-Myc tag at the 3’end of the MCM6 and 7 genes (Figure 1B) in Dm28c. The expressed tagged proteins were detectable in all cell lines (Figure 1A, 1B).

**Figure 1:**
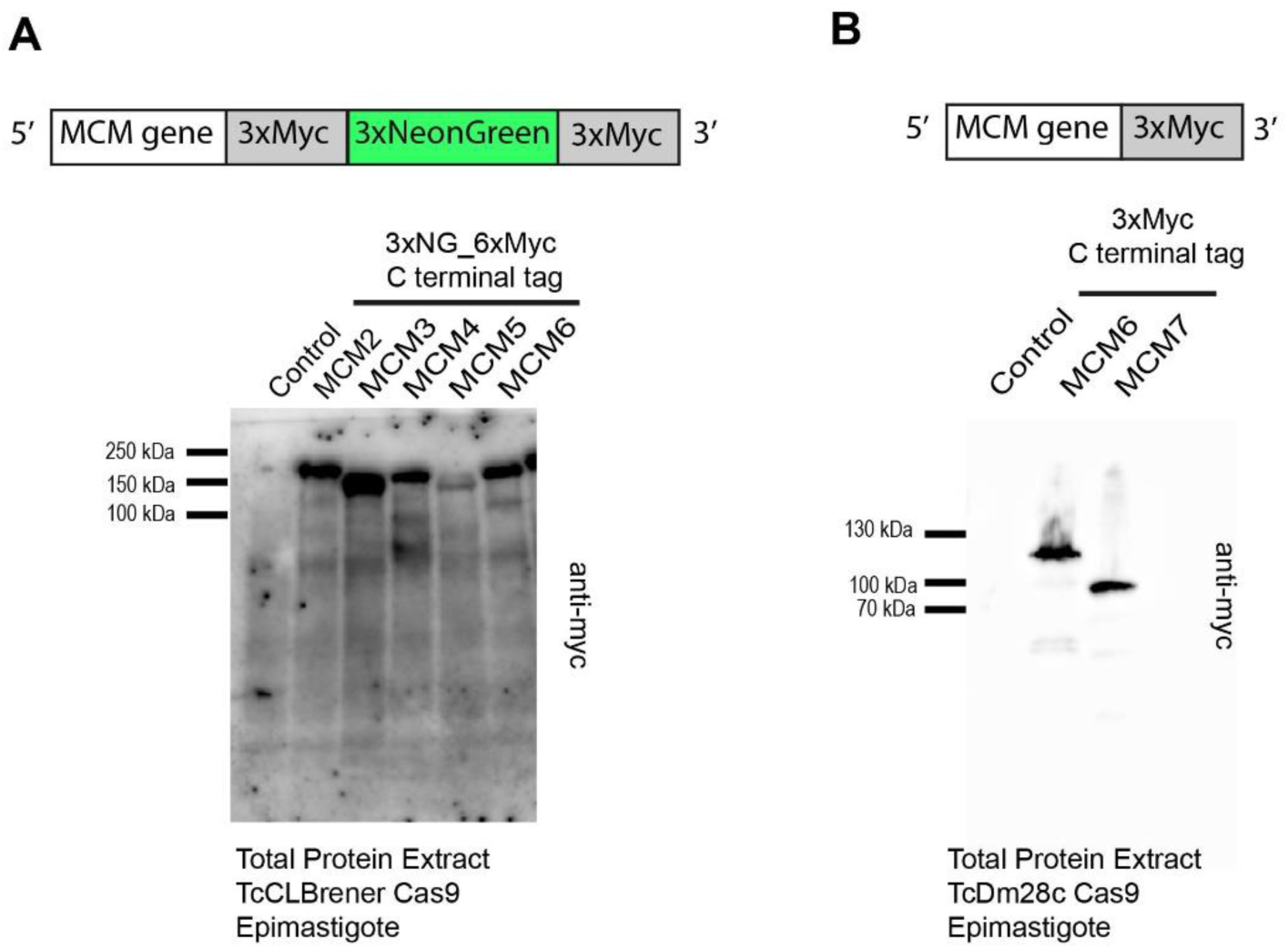
*T. cruzi* epimastigote cell lines modified by CRISPR/Cas9. **(A)** Western blot analysis of total protein from *T. cruzi* epimastigotes (CL Brener strain). As indicated in the illustration, three copies of mNeonGreen and six copies of the c-Myc epitope were inserted into the 3’ end of the MCM 2, 3, 4, 5, 6 genes. An anti-c-Myc antibody was used to identify the tagged proteins in total extracts. Protein extract of epimastigotes (TcCLBrener-Cas9) was used as a control. Expected band masses: ∼196 kDa for MCM2_Myc_NG; ∼178 kDa for MCM3_Myc_NG; ∼184 kDa for MCM4_Myc_NG; ∼172 kDa for MCM5_Myc_NG; ∼169 kDa for MCM6_Myc_NG. (B) Western blot analysis of total protein from *T. cruzi* epimastigote (Dm28c strain) cell lines. As indicated by the illustration, three copies of the c-Myc epitope were inserted into the 3’ end of the MCM 6 and 7 genes. An anti-c-Myc antibody was used to identify the tagged proteins in total extracts. Protein extract of epimastigotes (Dm28c-Cas9) was used as a control. Expected band masses: ∼103 kDa for MCM6_Myc and ∼86 kDa for MCM7_Myc

To determine the expression levels of MCM6 and MCM7 throughout the cell cycle, we synchronized *T. cruzi* Dm28c strain epimastigotes with hydroxyurea (Figure 2A). We obtained cell populations enriched at each cell cycle phase with the MCM6-Myc (Figure 2B) and MCM7-Myc modified cell lines (Figure 2E): 0h enriched for G1 and S phases, 4h enriched for S phase, 6h enriched for S and G2 phases, and 8h enriched for G2 phase. Whole protein extracts from these synchronized cells were used to assess the expression levels of MCM6-Myc (Figure 2C) and MCM7-Myc (Figure 2F) by western blotting, with FLAG-tagged Cas9 serving as the loading control. Quantification of band intensity from western blots of three independent replicates revealed that MCM6-Myc (Figure 2D) and MCM7-Myc (Figure 2G) are constitutively expressed throughout the epimastigote cell cycle with no significant variation among the phases.

**Figure 2:**
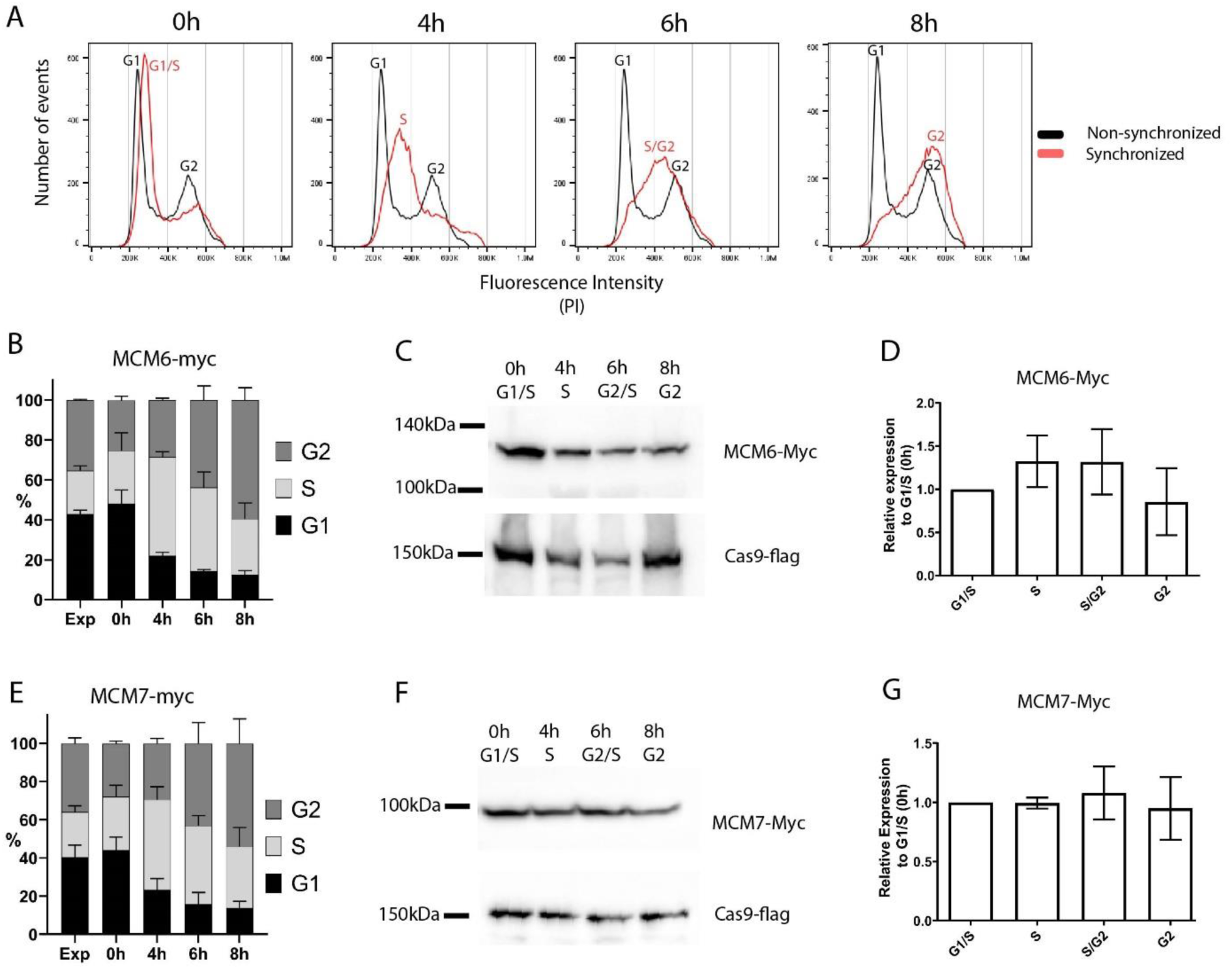
Cell cycle synchronization by hydroxyurea. *T. cruzi* epimastigotes from the Dm28c strain (MCM6-Myc and MCM7-Myc) were synchronized by hydroxyurea (HU) (see Methods for further details). **(A)** DNA content was measured by propidium iodide (PI) fluorescence intensity, using Flow Cytometry, and the histograms of number or events x PI intensity were analyzed in FlowJo software. The black line represents the cell cycle distribution of a non-synchronous population, and the two peaks for G1 and G2 populations are indicated (black letters). The red line represents the synchronous population at different time points after HU release (0h, 4h, 6h, 8h), and the cell cycle-enriched populations are marked on the peak histogram in red letters. **(B and E)** The histograms of number of events x PI intensity were used to determine the percentage of cells in each cell cycle stage (G1, S, G2) by FlowJo software, The graphs show this distribution at each time point of the synchronized and non-synchronized populations (B for MCM6-Myc cell line, E for MCM7-Myc cell line). **(C and F)** Western blot analysis of the synchronized cell population. Anti-Myc was used to detect **(C)** MCM6-Myc and **(F)** MCM7-Myc and anti-flag was used to detect the tagged Cas9 enzyme (input control). **(D and G)** Bands from the blot were quantified using Photoshop software, and the relative expression compared to the 0h point is represented in the graphs: **(D)** MCM6-Myc and **(G)** MCM7-Myc.

### MCMs are nuclear localised throughout the epimastigote cell cycle

We next investigated whether the cellular location of MCMs presents a different profile through the cell cycle, by performing immunofluorescence with clonal populations of the MCM-Myc tagged CL Brener (Supplementary Figure 1) and DM28c (Figure 3A) strains. Using morphological features such as the number of nuclei, kinetoplasts and flagella (Elias et al., 2007), we were able to determine the epimastigote cell cycle phase and location of MCMs. MCMs 2, 3, 4, 6 and 7 in CL Brener strain (Supplementary Figure 1) and MCM 6 (Figure 3A) and 7 (Figure 3B) in Dm28c strain were nuclear in all cell cycle phases, and no specific cytoplasmatic signal was identified. We also used the incorporation of the thymidine analogue EdU, to identify cells in S phase. Analysis of Z-stack images allowed us to reveal colocalization of MCM6-Myc, MCM7-Myc and EdU - regions visualized as yellow on merged images of anti-Myc (green) and EdU (red) staining (Figure 3C and D). As MCMs are part of the replisome (Polasek-Sedlackova et al., 2022), colocalization would be expected if MCM6 and MCM7 have a role in the DNA replication process.

**Figure 3:**
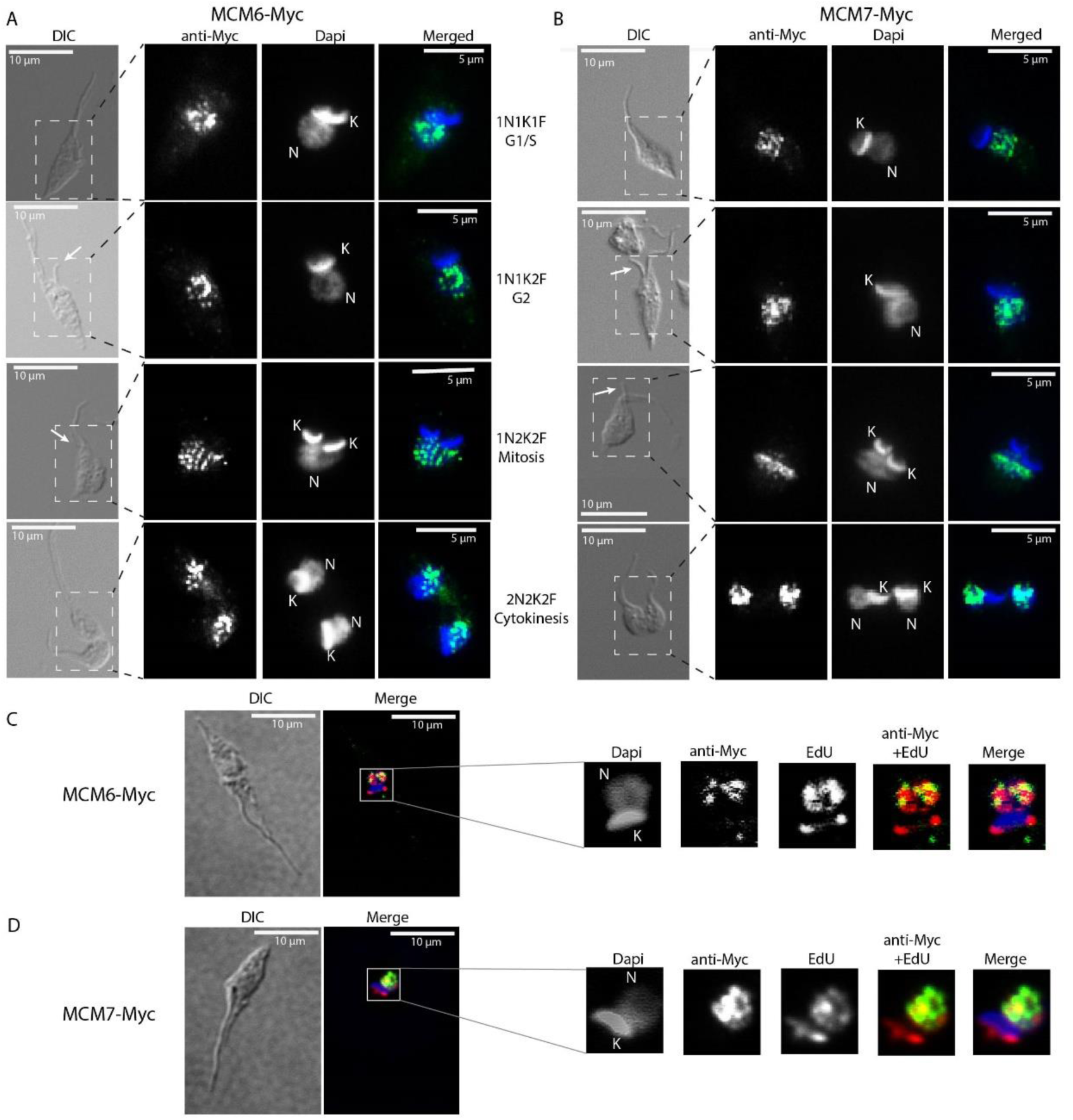
Immunofluorescence of MCM-tagged *T. cruzi* epimastigotes (Dm28c strain) using anti-Myc antibody. Epimastigotes from **(A and C)** the MCM6-Myc cell line (Dm28c strain) and **(B and D)** the MCM7-Myc cell line (Dm28c strain) were subjected to immunofluorescence using anti-Myc antibody (see Methods for further details). **(A and B)** Different cell cycle phases are represented and were identified by the N (Nucleus), K (Kinetoplast) and F (Flagellum) numbers. The white arrows indicate emerging new flagella. The dashed rectangle in the DIC images highlight the regions shown in the corresponding fluorescent images (ani-Myc, DAPi and merged). **(C and D)** Epimastigotes in S phase of the cell cycle were identified by EdU incorporation (red). The yellow areas are sites of colocalization of (C) MCM6-Myc and EdU, and (D) MCM7-Myc and EdU. A and B images were acquired on Microscope Olympus BX51. C and D were acquired on Microscope Olympus IX81 and the images are a layer from a Z-stack acquisition, deconvoluted by software Auto Quant. White bar scale (10 µm), Green (anti-Myc signal, yellow when on a red background), Blue (DAPI signal), Red (EdU signal), DIC (Differential Interference Contrast).

### MCM6-Myc and MCM7-Myc cells are arrested in G1 phase during metacyclogenesis

During the *T. cruzi* lifecycle, replicative epimastigotes differentiate into metacyclic trypomastigotes, which are infective but non-replicative. This differentiation process, known as metacyclogenesis, occurs within the vector gut (Ferreira et al., 2023) and can also be induced *in vitro* (Contreras et al., 1985) (Figure 4A). We induced *in vitro* metacyclogenesis and collected samples at four-time points to analyze the cell cycle and the expression of MCM6-Myc and MCM7-Myc. Epimastigotes were cultured until they reached stationary growth phase (point 2 in Figure 4A), and then subjected to stress in TAU medium for 2 h (point 3 in Figure 4A). Subsequently, they were incubated for 5 days in TAU3aaG medium (point 4 in Figure 4A) to allow metacyclogenesis. As a control, epimastigotes from the stationary phase were diluted in fresh medium and collected 24h later, representing replicative epimastigotes recovered from a non-replicative state. At the end of metacyclogenesis, we quantified the percentage of cells that had completely differentiated into metacyclic trypomastigotes, as well as intermediate forms and non-differentiated epimastigotes (Figure 4B). Cellular morphology, including the shape and positioning of the nucleus and kinetoplast, was used to identify epimastigotes, intermediates, and metacyclic trypomastigotes (Gonçalves et al., 2018). For the MCM6-Myc cell line, 37.5% of the cells had completely differentiated into metacyclic trypomastigotes, while for the MCM7-Myc cell line, 32.7% of the cells were metacyclic trypomastigotes (Figure 4B).

**Figure 4:**
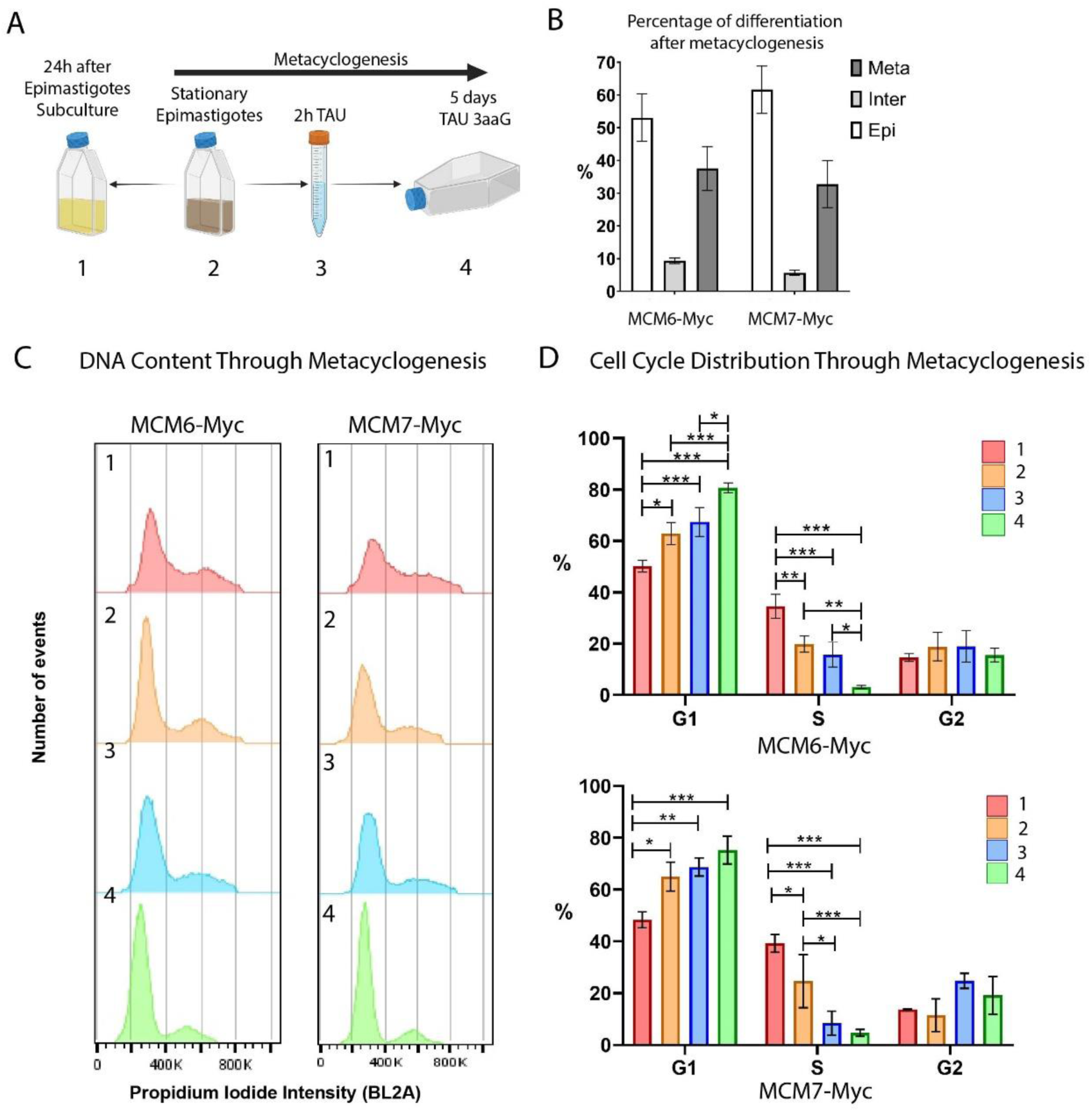
Metacyclogenesis of MCM6-Myc and MCM7-Myc cell lines (Dm28c strain). **(A)** Schematic representation of the metacyclogenesis process *in vitro*. (1) Epimastigotes in the stationary phase of growth were diluted in fresh medium and collected 24h later, serving as the control for replicating epimastigotes. (2) Epimastigotes in the stationary phase were (3) stressed for 2 h in TAU medium and (4) differentiated over 5 days in TAU 3aaG medium (see Methods for further details). (B) After metacyclogenesis, cells were fixed on slides and stained with DAPI. Epimastigotes, intermediate forms, and metacyclic trypomastigotes were counted. The bar chart shows the percentage of each cell type, averaged from three replicates. (C) The DNA content of each sample collected during metacyclogenesis was analyzed by flow cytometry. The histograms represent the number of events versus propidium iodide intensity (BL2A) for each metacyclogenesis time point. (D) The histograms from (C) were analyzed using FlowJo software, and cell cycle phases were determined by Dean-Jett-Fox cell cycle modeling. The bar charts show the percentage of parasites in each cell cycle phase throughout metacyclogenesis, averaged from three replicates. A two-way ANOVA was applied, and significance is indicated by * (p≤0.01), ** (p≤0.001), and *** (p≤0.0001).

Using flow cytometry, we analyzed the DNA content by propidium iodide (PI) intensity for each time point during metacyclogenesis (Figure 4C). We then quantified the cell cycle distribution using the cell cycle modeling tool from FlowJo software (Figure 4D).

In both cell lines, MCM6-Myc and MCM7-Myc, we observed that the percentage of cells in G1 phase increased as epimastigotes transitioned from the replicative phase (point 1) to the stationary growth phase (point 2), reaching a maximum at the end of metacyclogenesis (Figure 4D). In the MCM6-Myc cell line, the percentage of cells in G1 phase in replicative epimastigotes (point 1) was 50.2%, compared to 62.9% in epimastigotes in the stationary growth phase (point 2). At the end of metacyclogenesis, (point 4) the percentage of cells in G1 was 80.7% (Figure 4D, upper graph). There was a similar trend with MCM7-Myc cell line, with 48.3% in G1 phase in replicative epimastigotes (point 1), 65% in epimastigotes in the stationary growth phase (point 2), and 75.3% at the end of metacyclogenesis (point 4 in Figure 4D, lower graph).

Simultaneously, in both cell lines, the percentage of parasites in S phase gradually decreased during the transition from replicative epimastigotes to non-replicative metacyclics. In the MCM6-Myc cell line, the percentage of epimastigotes in S phase (point 1) decreased from 34.6%, to 19.9% in the stationary growth phase (point 2). At the end of metacyclogenesis (point 4), the percentage of parasites in S phase was 3.1% (Figure 4D, upper graph). In the MCM7-Myc cell line, the percentage of epimastigotes in S phase (point 1) was 39.2%, compared to 24.6% in the stationary growth phase (point 2). At the end of metacyclogenesis (point 4), the percentage of cells in the S phase was 4.6% (Figure 4D, lower graph). Finally, in the tagged lines, there was no significant difference in the percentage of parasites in G2 at the 4 points collected during metacyclogenesis.

### Expression levels of MCM6-Myc and MCM7-Myc decrease through metacyclogenesis

Given that cells exiting the cell cycle exhibit decreased levels of MCM (Madine et al., 2000) and that we have previously observed that metacyclic trypomastigotes do not express MCM7 (Calderano et al., 2014), we investigated the expression profiles of MCM6-Myc and MCM7-Myc at four distinct points during metacyclogenesis (Figure 5A). Our observations revealed a decrease in expression levels in both parasite lines, from replicative epimastigotes (point 1) to parasites post-metacyclogenesis (point 4, Figure 5A).

**Figure 5:**
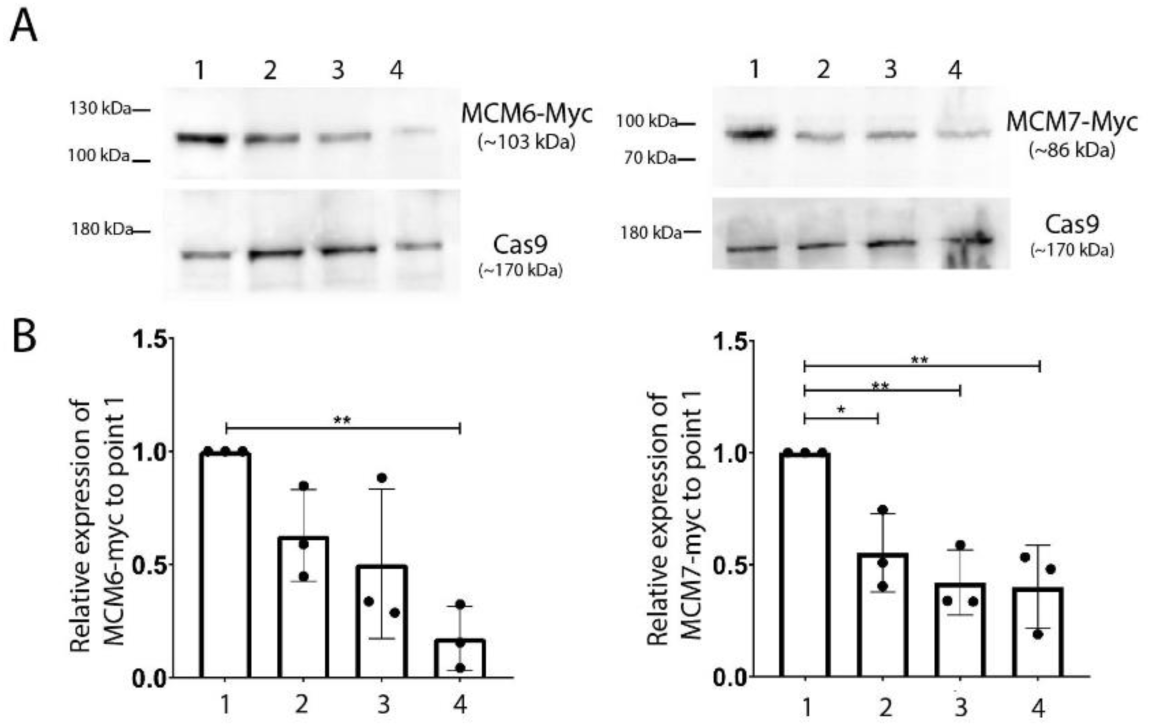
Western blotting analysis of MCM6-Myc and MCM7-Myc during metacyclogenesis of *T. cruzi* (Dm28c strain). **(A)** Whole protein extracts from each metacyclogenesis point were subjected to western blot analysis. An anti-Myc antibody was used to identify MCM6-Myc and MCM7-Myc, while an anti-Flag antibody was used to detect Cas9 expression, serving as an input control. **(B)** The bands detected in **(A)** were quantified using Photoshop software. The bar chart shows the expression levels relative to replicative epimastigotes (point 1), averaged from 3 replicates. One-way ANOVA test was applied, and significance is indicated by * (p<0.05) and ** (p<0.01).

To quantify these changes, we measured band intensity from three independent Western blot replicates and analyzed the MCM expression levels at each point (points 1 to 4, Figure 5A) relative to the levels in replicative epimastigotes (point 1, Figure 5A). In the MCM6-Myc cell line, the expression levels in stationary epimastigotes (point 2), and in epimastigotes after 2 h in TAU (point 3), followed a reducing trend compared to those in replicative epimastigotes (Figure 5B). The reduction in expression levels was more pronounced and statistically significant in parasites post-metacyclogenesis (point 4). In the MCM7-Myc cell line, a similar pattern was observed. The decreased expression levels from replicative epimastigotes (point 1, right graph, Figure 5B) to stationary epimastigotes (point 2), 2 h TAU-stressed epimastigotes (point 3), and parasites post-metacyclogenesis (point 4) were statistically significant across all points (Figure 5B).

### MCM6-Myc and MCM7-Myc are nuclear-localized in amastigotes and stationary-epimastigotes, but are not expressed in metacyclic or tissue-cultured trypomastigotes

MCMs are nuclear-localised throughout the entire cell cycle in replicative epimastigotes (Figure 3). Given that expression of MCM6-Myc and MCM7-Myc diminishes in stationary epimastigotes, we investigated the impact on their cellular location. To confirm that epimastigotes in the “stationary growth phase” are non-replicating, we assessed their proliferation capacity using EdU incorporation, followed by analysis via flow cytometry (Figure 6A and 6B). The results showed that replicating epimastigotes incorporated EdU, whereas stationary epimastigotes did not (Figure 6A, 6B). When we analyzed the cellular location of MCM6-Myc and MCM7-Myc, we found that they had the same nuclear pattern as in stationary-epimastigotes (Figure 6 C and D). Additionally, after metacyclogenesis process, in epimastigotes that did not differentiate, MCM6-Myc and MCM7-Myc were also nuclear-localised (Supplementary Figure 3). However, no specific signal was detectable in fully differentiated metacyclic trypomastigotes (Figure 7 A).

**Figure 6:**
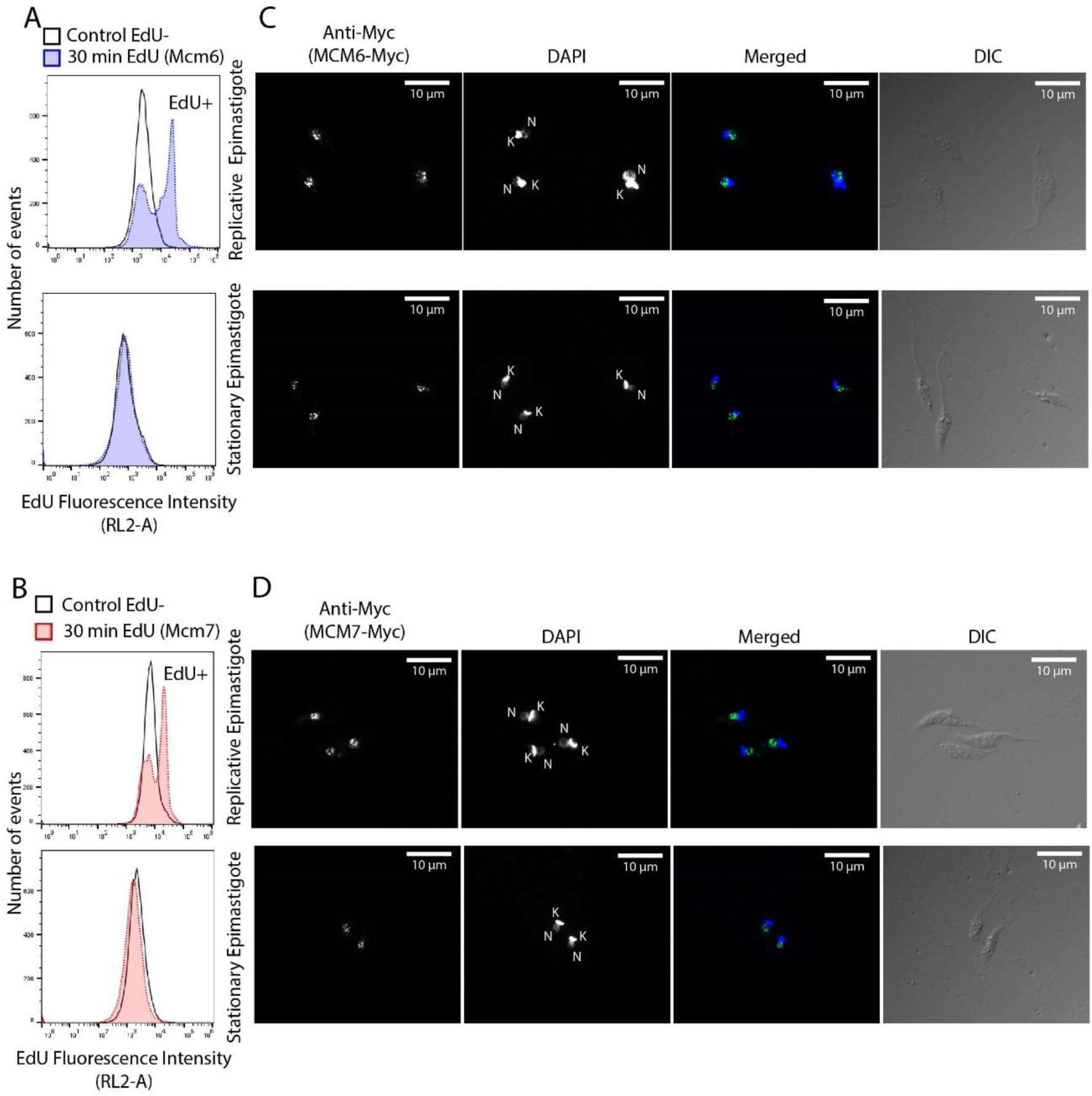
Cellular localization of MCM6-Myc and MCM7-Myc in replicative and non-replicative *T. cruzi* epimastigotes (Dm28c strain). *T. cruzi* epimastigotes in log phase (replicative) and stationary phase (non-replicative) were analyzed to determine MCM localization **(A and B)** Cells were incubated with EdU for 30 minutes (see Methods for further details) and analyzed by flow cytometry. The histograms show the number of cells versus EdU fluorescence intensity. Blank histograms represent the negative control, where parasites (MCM6-Myc cell line and MCM7-Myc cell line) were not incubated with EdU. In **(A)**, the blue histograms represent the MCM6-Myc line incubated with EdU. In **(B)**, the red histograms represent the MCM7-Myc line incubated with EdU. **(C and D)** Immunofluorescence of epimastigotes using an anti-Myc antibody to detect **(C)** MCM6-Myc and **(D)** MCM7-Myc. White bar scale represents 10 µm. N, nucleus; K, kinetoplast; green, anti-Myc signal; blue, DAPI (DNA); DIC, Differential Interference Contrast.

**Figure 7:**
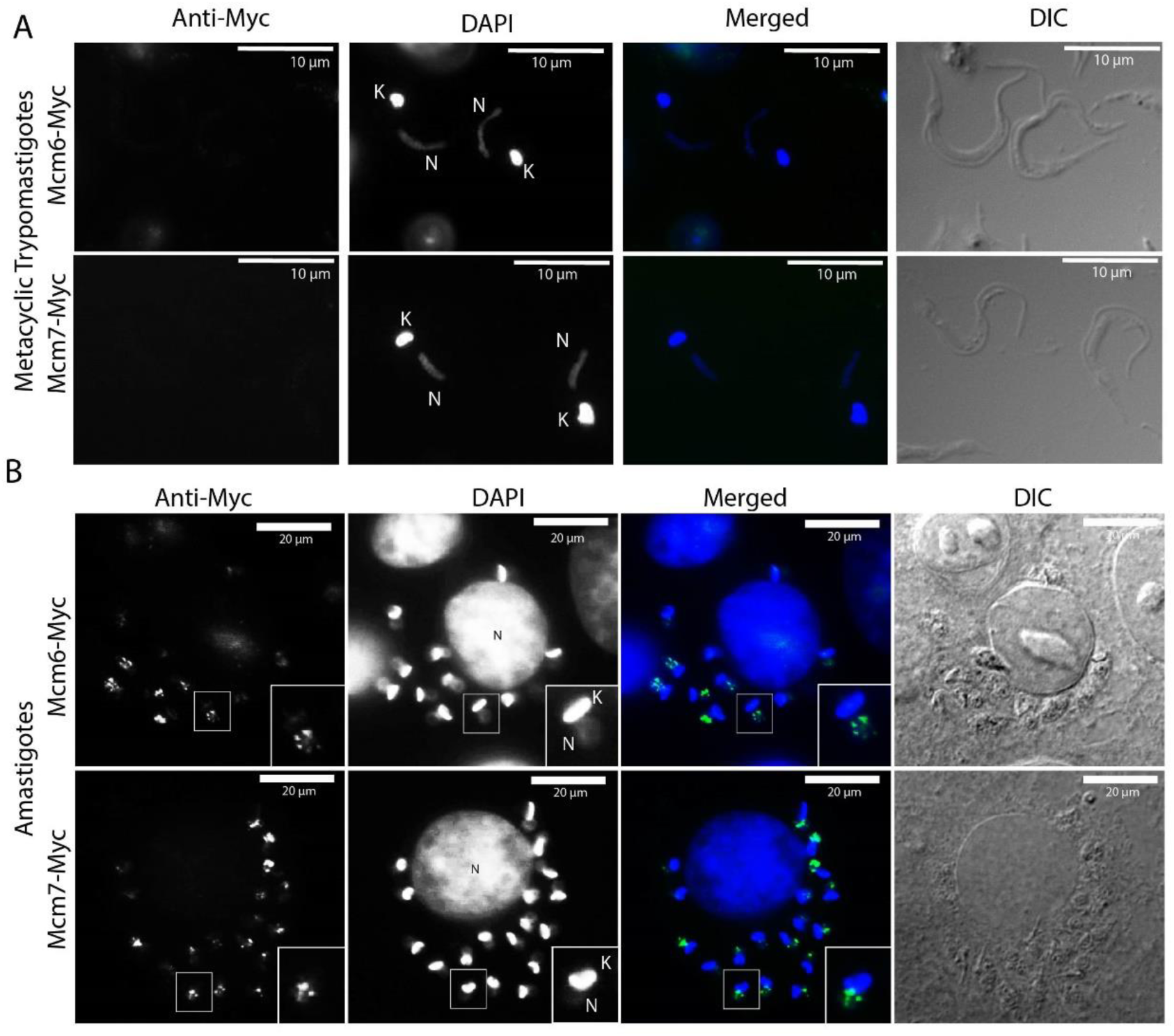
Immunofluorescence of *T. cruzi* metacyclic trypomastigotes and amastigotes (Dm28c strain). Metacyclic trypomastigotes (in **A**) and intracellular amastigotes (in **B**) derived from the MCM6-Myc (Dm28c strain) and MCM7-Myc cell lines (Dm28c strain) were subjected to immunofluorescence using anti-Myc antibody. White bar scale represents 10 µm in A and 20 µm in B. N, nucleus; K, kinetoplast; green, anti-Myc; blue, DAPI (DNA); DIC, Differential Interference Contrast. In B, the image corresponds to one slice of a Z-stack image, and the framed amastigote in the image is magnified twofold in the bottom right corner of anti-Myc, DAPI, and merged images.

Metacyclic trypomastigotes were then used to infect mammalian cells to produce amastigotes and TCTs (Tissue-Culture Trypomastigotes). Using immunofluorescence to assess MCM6-Myc and MCM7-Myc locations within intracellular amastigotes, we observed that these proteins were also confined to the nucleus in this replicative life cycle stage (Figure 7B). Unlike epimastigotes, the cell cycle stage of amastigotes cannot be inferred from morphology. However, no cytoplasmic signal was detected for either MCM protein, with localization being exclusively nuclear.

### Tissue-culture trypomastigotes are arrested in G1 phase of the cell cycle

Given our previous observation that cells are arrested in the G1 phase following metacyclogenesis, we characterized the cell cycle profile of TCTs using the Dm28c MCM6-Myc and MCM7-Myc strains, with cell cycle stages defined using the FlowJo software modeling tool. The cell cycle profiles of replicative epimastigotes and TCTs of MCM6-Myc and MCM7-Myc cell lines were compared (Figure 8A and B, respectively). ∼50% of epimastigotes were in G1 phase (Figure 8C and D), 50.2% and 48.3% in the MCM6-Myc and MCM7-Myc cell lines, respectively. By comparison, TCTs exhibited a higher proportion, with around ∼74% of cells in G1 phase (Figure 8C and D). Therefore, G1 arrest was also observed in TCTs. We performed immunofluorescence on both epimastigotes and TCTs to further assess MCM protein expression. As shown (Figure 8E), specific nuclear signals were observed exclusively in epimastigotes, while TCTs did not exhibit expression of MCM6-Myc or MCM7-Myc.

**Figure 8:**
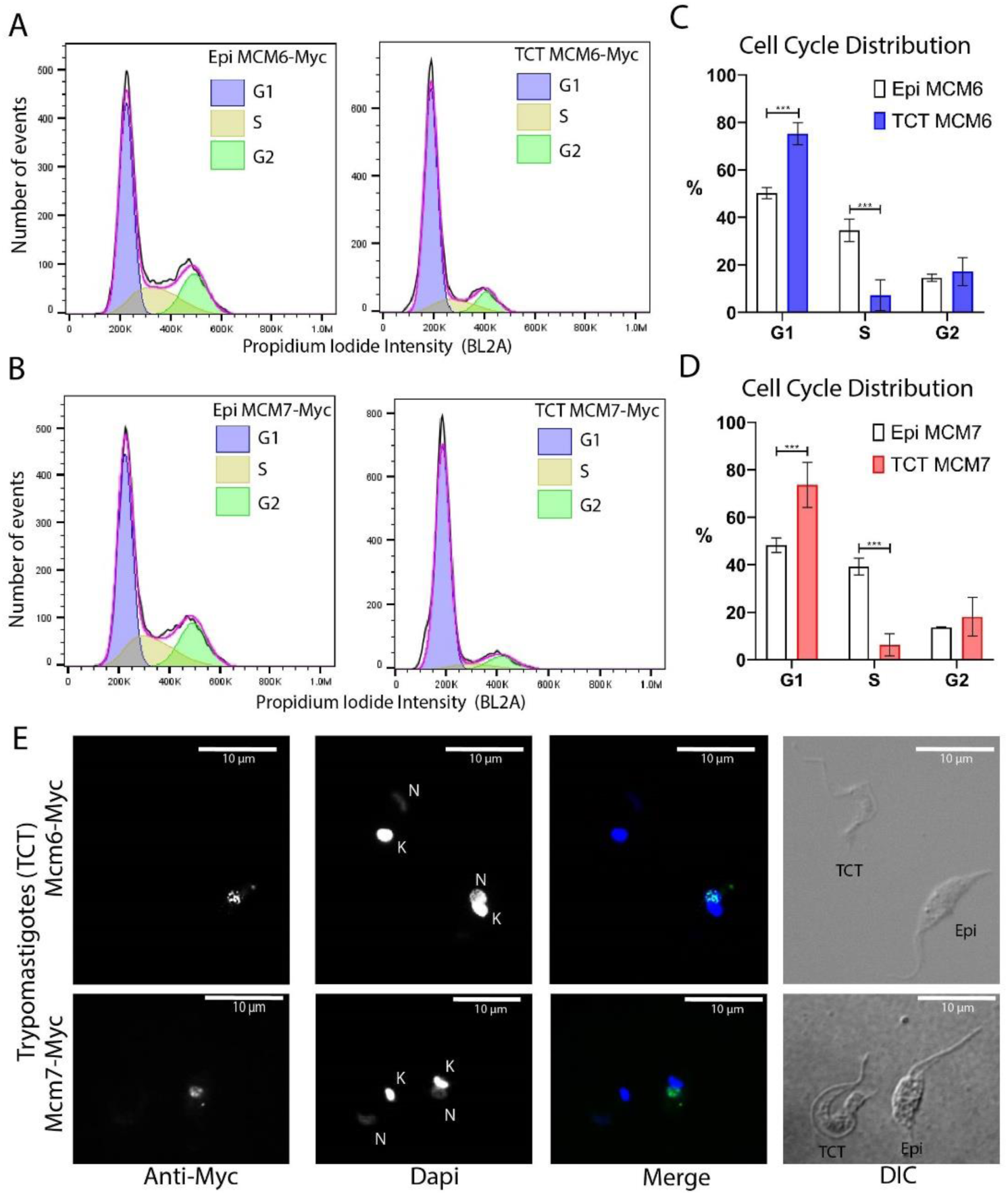
Cell cycle distribution and immunofluorescence analysis of tissue culture trypomastigotes (Dm28c strain). (**A** and **B**) Histograms of event counts versus propidium iodide intensity from flow cytometry analysis of epimastigotes and TCTs. Cell cycle distribution was determined using the cell cycle modeling tool in FlowJo software. (**A**) MCM6-Myc cell line. (**B**) MCM7-Myc cell line. (**C** and **D**) Bar charts illustrating the cell cycle distribution of epimastigotes and TCTs. (**C**) MCM6-Myc cell line. (**D**) MCM7-Myc cell line. (**E**) Immunofluorescence analysis of TCTs and epimastigotes on the same slide, using an anti-Myc antibody in the MCM6-Myc and MCM7-Myc cell lines. The white bar scale represents 10 µm. White bar scale represents 10 µm. N, nucleus; K, kinetoplast; green, anti-Myc; blue, DAPI (DNA); DIC, Differential Interference Contrast.

## Discussion

The MCM2-7 complex is the only pre-RC component that is integrated into the replisome during DNA replication (Polasek-Sedlackova et al., 2022). In *T. cruzi,* MCM expression is exclusive to the replicative forms (Calderano et al., 2014) (Figures 3 and 7 and Supplementary Figure 1), localizes inside the nucleus (Figure 3A and B and Supplementary Figure 1) and colocalizes with the site of EdU incorporation (Figure 3C and D), supporting the view that MCM has a conserved role in DNA replication and is part of the replisome in this organism.

The pre-RC complex demarcates the DNA replication starting sites that must be fired only once per cell cycle, so their components are subject to three main regulation mechanisms: degradation, nuclear export, and chromatin detachment. Generally in eukaryotes, MCM subunits are nuclear throughout the cell cycle and detached from chromatin at the end of the S phase (Forsburg, 2004). Exceptionally, they can be exported from the nucleus as observed in *Saccharomyces cerevisiae* (Dalton and Whitbread, 1995). When cells exit the cell cycle, MCMs are downregulated and expression is restricted to low levels in quiescent cells (Blow and Hodgson, 2002, Stoeber et al., 2001). However, in terminally differentiated cells MCM expression is abolished (Carroll et al., 2018). In *T. cruzi,* we found that MCMs are expressed and nuclear-localized throughout the epimastigote cell cycle (Figures 2 and 3, Supplementary Figure 1), in line with most eukaryotes (Forsburg, 2004), including *T. brucei* (Dang and Li, 2011). Variation in MCM expression levels was only observed during the differentiation from epimastigotes to metacyclic trypomastigotes, where MCMs were downregulated (Calderano et al., 2014) (Figures 4D and 5). The first step for *in vitro* metacyclogenesis is culturing epimastigotes until the stationary phase, which is pre-adaptative to the differentiation process (Hernández et al., 2012). It is characterized by higher rates of protein degradation (Henriquez et al., 1993) and changes in metabolism caused by nutrientscarcity (Barison et al., 2017). These stationary epimastigotes are G1 arrested-they do not replicate (Figure 6) and are enriched in the G1 phase (Figure 4D)—and exhibit a slight decrease in MCM expression, significant only in the case of MCM7 (Figure 5). Hence, maintaining MCM expression in these arrested cells guarantees their readiness to re-enter the cell cycle and initiate DNA replication promptly when favorable conditions are restored (Lee and Osley, 2021). The MCM downregulation was more pronounced at the final stage of metacyclogenesis (Figure 5). While differentiated metacyclic trypomastigotes do not express MCMs (Figure 7A and Supplementary Figure 3), the remaining epimastigotes in the culture (Figure 4 B) still express MCMs that are restricted to the nucleus (Supplementary Figure 3). This nuclear pattern of MCM expression in epimastigotes was consistent in all analyzed conditions, which were replicating (Figure 3 and Supplementary Figure 1), stationary (Figure 6) and committed to the differentiation process (Supplementary Figure 3). Therefore, mechanisms other than nuclear export and degradation may be involved in regulating MCM activity in epimastigotes. MCM down-regulation was only observed after metacyclogenesis, where MCM6 and 7 were completely depleted in differentiated metacyclic trypomastigotes (Calderano et al., 2014) (Figure 7). Protein degradation rates start to increase in stationary epimastigotes (Henriquez et al., 1993), and higher levels of turn-over occur during metacyclogenesis, where autophagy and proteasome-mediated protein degradation play a crucial role, and their inhibition impairs this differentiation process (Losinno et al., 2020, Cardoso et al., 2011, Cardoso et al., 2008). Therefore, MCM degradation represents a potential additional regulatory mechanism that acts to inhibit DNA replication in differentiated metacyclic trypomastigotes (Calderano et al., 2014). Different proteomic approaches have identified proteins differentially expressed in stationary epimastigotes (Avila et al., 2018) and during metacyclogenesis (Amorim et al., 2017, Lucena et al., 2019), however, MCMs were not detected in these studies. Detection of MCMs by proteomics seems to be challenging; studies investigating the cell cycle (Chávez et al., 2021), nuclear (dos Santos Júnior Ade et al., 2015) and chromatin (Leandro de Jesus et al., 2017) proteome profiles have been unable to detect these proteins, with the exception of MCM2 (TcCLB.506933.40). This protein was detected in the proteome of the two cell cycle phases analyzed, G1 and S (Chávez et al., 2021), consistent with the constitutive expression of MCM 6 and 7 reported here (Figure 2 C and F). With this exception, the limited ability to date of proteome techniques to detect MCMs prevents more in-depth comparisons between our data and previous studies.

During the transition from a replicative to a non-replicative state, the percentage of cells in G1 progressively increased (Figure 4 D), consistent with findings by Santos et al., 2018. This increase in G1-phase cells peaked at the end of metacyclogenesis (Figure 4D) and a similar G1 arrest was observed here with non-replicative TCTs (Figure 7 A-D).). Although DNA content analysis cannot differentiate between G1 and G0 arrested cells due to their identical DNA content, these populations are distinct and exhibit different transcriptome (Coller et al., 2006) and proteome (Ly et al., 2015) profiles. Several markers for G0 cells have been identified (Breeden and Tsukiyama, 2022), including unlicensed origins of replication (Carroll et al., 2018, Stoeber et al., 1998), reduced rRNA transcription (Hannan et al., 2000), decreased translation (Pereira et al., 2015; Liu and Sabatini, 2020), diminished transcription rates (Choder, 1991, Young et al., 2017, McKnight et al., 2015, Mews et al., 2014), and condensed chromatin (Evertts et al., 2013, Rawlings et al., 2011, Schmiady and Sperling, 1981, Swygert et al., 2019). Additionally, *S. cerevisiae* in the stationary phase are heterogeneous and consist of quiescent (G0) and non-quiescent cells (Allen et al., 2006, Aragon et al., 2008). In this context, we categorize stationary epimastigotes as being in a G1/G0 arrested state, as they are enriched in G1 (Figure 4D) and do not synthesize DNA (Figure 6). We considered the non-replicative lifeforms trypomastigotes (both metacyclic and TCT forms) to be in G0 arrest, as they exhibit several hallmarks of G0 cells, including reduced transcriptional (Elias et al., 2001) and translational (Smircich et al., 2015) activities, increased chromatin condensation (Lima et al., 2022), absence of a nucleolus (Lima et al., 2022), and unlicensed DNA replication origins. This latter feature is supported by the lack of MCM6 and MCM7 expression and the detachment of Orc1/Cdc6 from chromatin (Calderano et al., 2014).

G1 accumulation during metacyclogenesis suggests that the exit from the cell cycle occurs at the G1 phase, with differentiated trypomastigotes arrested in G0. Notably, in *T. brucei*, the transition from the replicative slender form to the non-replicative stumpy form occurs in early G1(Briggs et al., 2021), raising the possibility of a G1 checkpoint that governs cell fate in these trypanosomes, as is well established in other eukaryotes (Johnson and Skotheim, 2013). In *T. cruzi*, G1 arrest is also observed in intracellular amastigotes following stress induction (Dumoulin and Burleigh, 2018), with the cell cycle resuming after stress removal. In *S. cerevisiae*, nutrient availability and cell size are the primary factors required to pass the G1 restriction point (Johnston et al., 1977, Johnson and Skotheim, 2013). When nutrient levels are sufficient, cells grow to the appropriate size and then commit to cell cycle progression. In *T. cruzi*, it is well-established that nutritional starvation is a key trigger for metacyclogenesis and consequent exit from the cell cycle (Gonzales-Perdomo et al., 1988, Figueiredo et al., 2000, Hamedi et al., 2015, Barison et al., 2017). Moreover, recent research has shown that *T. cruz*i epimastigotes undergo asymmetric cell division, with the smaller daughter cell exhibiting a prolonged G1 phase, indicating that a minimum cell size must be achieved before entering S phase (Campbell and de Graffenried, 2023). These observations, together with the G1 arrest seen after metacyclogenesis and in TCTs, suggest the presence of a G1 restriction point in *T. cruzi*. The G1 checkpoint is conserved among eukaryotes, with the exception of *Schizosaccharomyces pombe*, and is typically regulated by transcriptional control (Johnson and Skotheim, 2013). In both unicellular and multicellular organisms, a transcription factor is bound to a repressor that, when phosphorylated by cyclin-dependent kinases (CDKs), releases the transcription factor to activate the expression of genes involved in cell cycle progression. This repression of transcription factors is maintained when cells exit the cell cycle. However, in trypanosomatids, transcription is polycistronic (Imboden et al., 1987), meaning they cannot regulate transcription via RNA polymerase II as in other eukaryotes (Clayton, 2019). As a result, the mechanisms governing the G1-G0 transition in these parasites remain unclear. Nonetheless, it is intriguing that this G1 control checkpoint appears to be present in this early-branching eukaryote, which may also employ alternative regulatory mechanisms beyond transcriptional control.

In conclusion, epimastigotes arrested in G1/G0 can either maintain MCM expression and resume the cell cycle when conditions become favorable or undergo differentiation, entering G0 as MCMs are degraded as part of the replication repression mechanism. However, the regulatory mechanisms governing this process remain unclear and warrant further investigation.

## Acknowledgments

We thank Dr. Noelia Lander and Dr. Roberto Docampo for pMOTag23M plasmid and FAPESP for grant and fellowship funding.

## Author Contributions

Conceptualization: SGC, MCE. Methodology: FCC, MCT, BAS, APC, EM. Investigation: SGC, BAS, APC, EM. Visualization: SGC, BAS, APC, EM. Funding acquisition: SGC, MCE, JMK. Project administration: SGC. Supervision: JMK, SGC. Writing – original draft: SGC with comments from authors. Writing – review & editing: SGC, MCE, JMK, MCT, FCC, BAS, APC, EM.

## Supplementary Figures

**Supplementary Figure 1:**
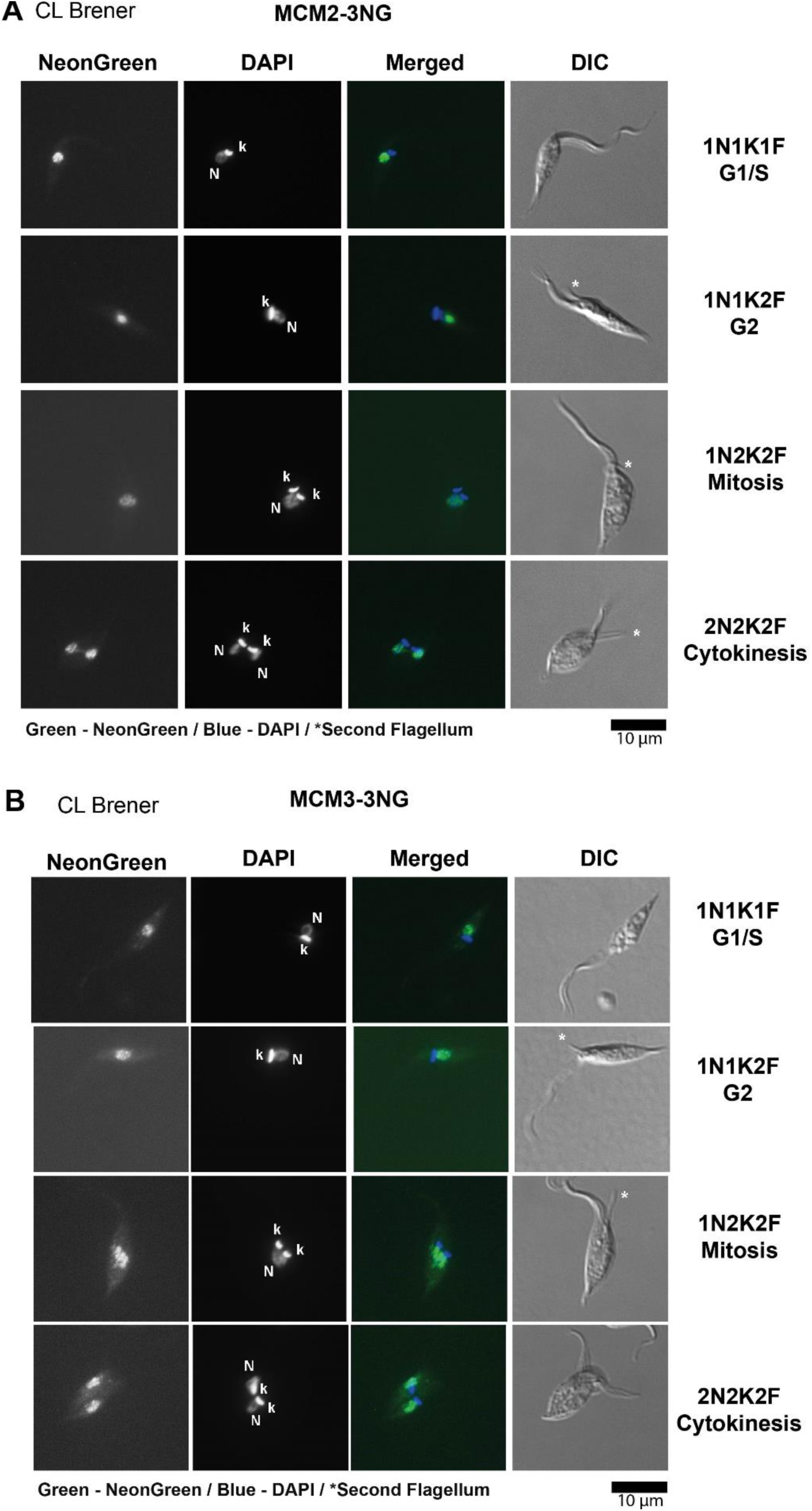
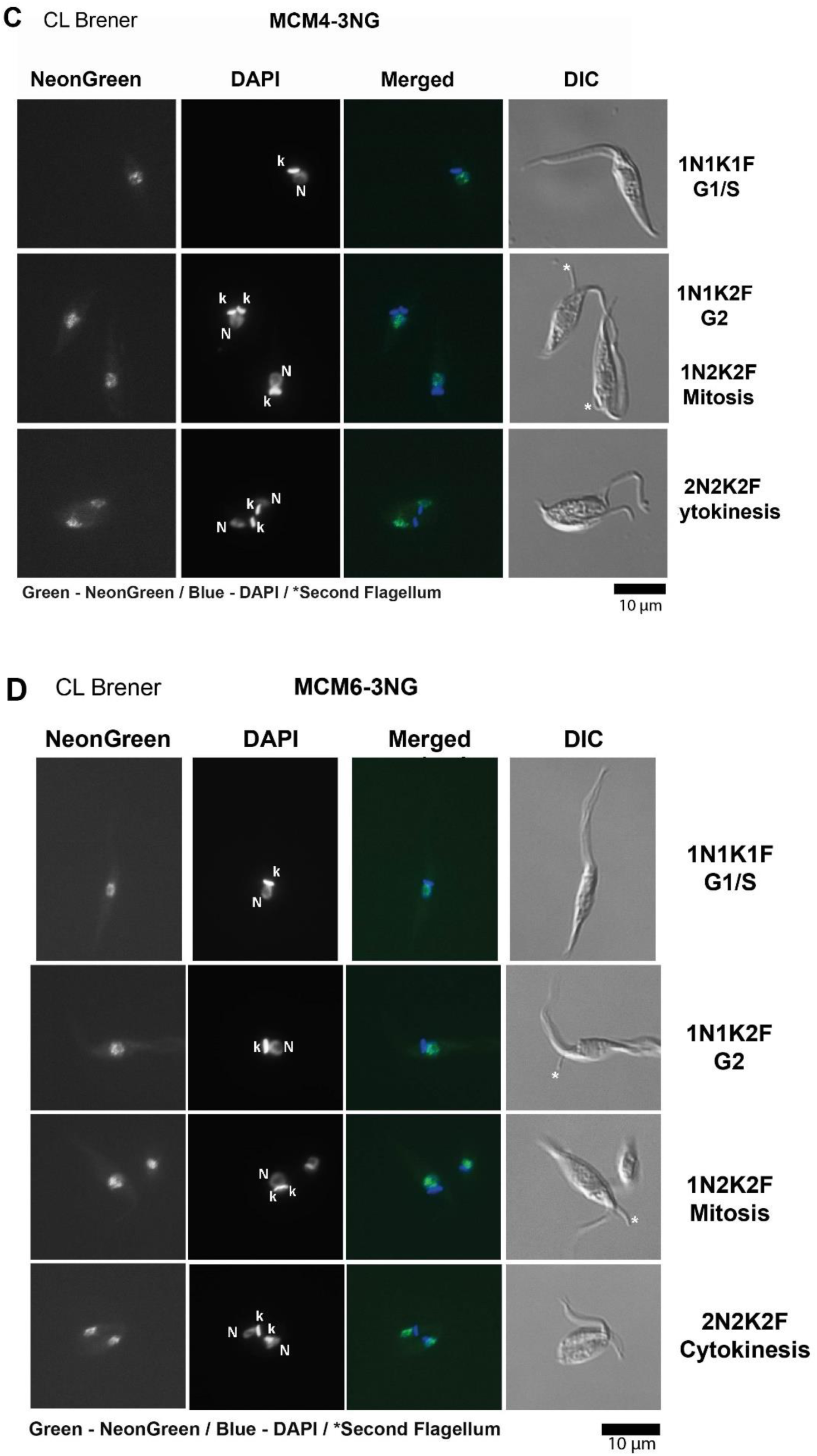
Fluorescence imaging of *T. cruzi* CL Brener strain epimastigotes modified using CRISPR/Cas9. *T. cruzi* epimastigotes were genetically modified by CRISPR/Cas9 to incorporate three copies of the mNeonGreen gene at the 3’ end of the MCM2, MCM3, MCM4, and MCM6 genes. Fluorescence microscopy was employed to capture images of mNeonGreen fluorescence (green), DAPI-stained DNA (blue), and Differential Interference Contrast (DIC). Cells at different stages of the cell cycle are shown. The black scale bar represents 10 µm. N indicates the nucleus, K indicates the kinetoplast, and F indicates the flagellum. (A) MCM2-3NG cell line; (B) MCM3-3NG cell line; (C) MCM4-3NG cell line; (D) MCM6-3NG cell line.

**Supplementary Figure 2:**
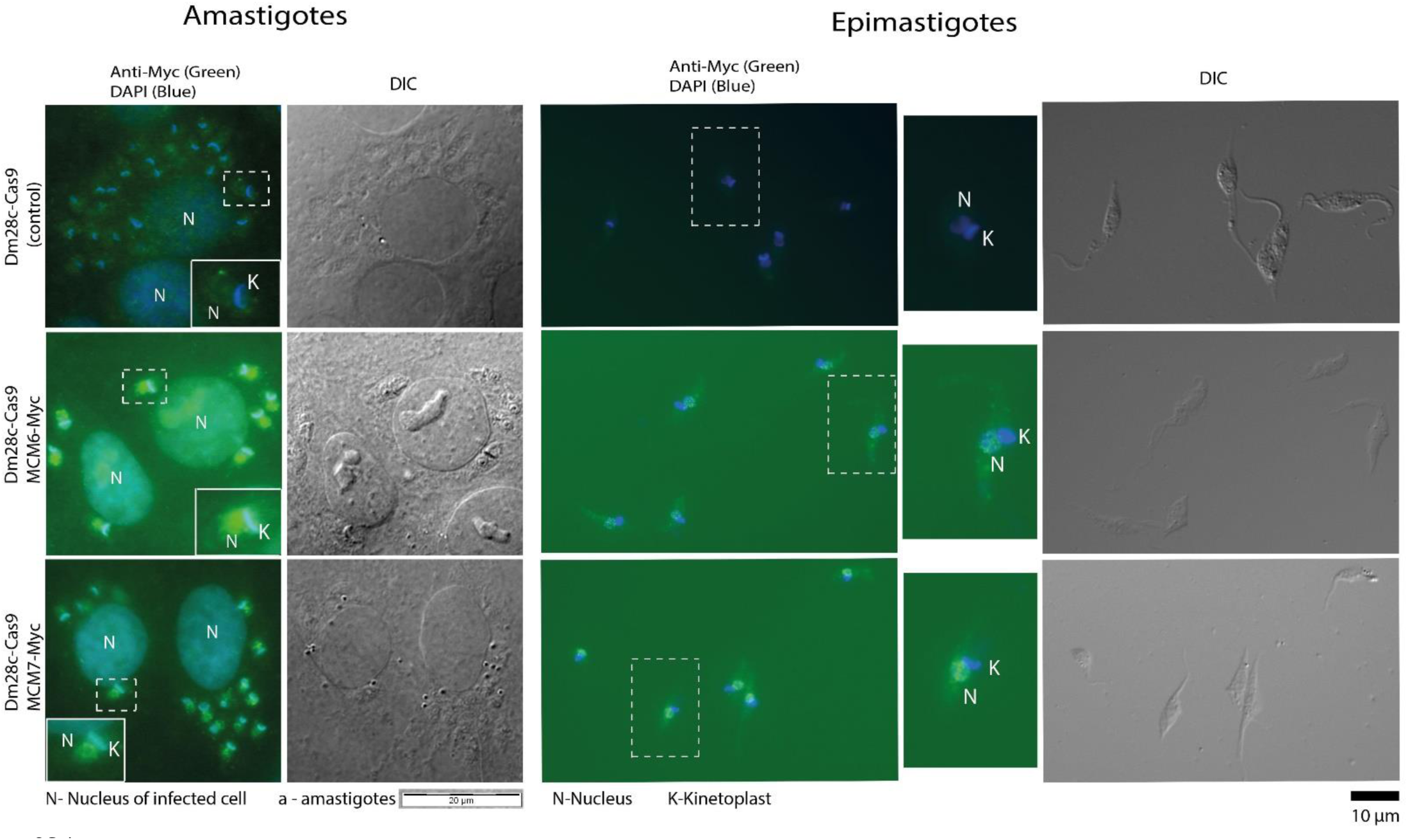
Immunofluorescence imaging of *T. cruzi* Dm28c strain epimastigotes and amastigotes genetically modified using CRISPR/Cas9. The epimastigotes were modified by inserting three copies of the Myc sequence at the 3’ end of the MCM6 and MCM7 genes. Immunofluorescence microscopy was performed to visualize anti-Myc staining (green), DAPI-stained DNA (blue), and Differential Interference Contrast (DIC) images. Here are shown raw images of intracellular amastigotes and epimastigotes from: Dm28c-Cas9 (control), Dm28c-MCM6-Myc, and Dm28c-MCM7-Myc cell lines. The black scale bar represents 10 µm, and the white scale bar represents 20 µm. Dashed rectangles highlight cells that are 2x magnified. For amastigotes, magnified images are on the bottom right (control and MCM6-Myc) and bottom left (MCM7-Myc). For epimastigotes, the magnified cells are located to the left of the immunofluorescence image. N denotes the nucleus and K the kinetoplast.

**Supplementary Figure 3:**
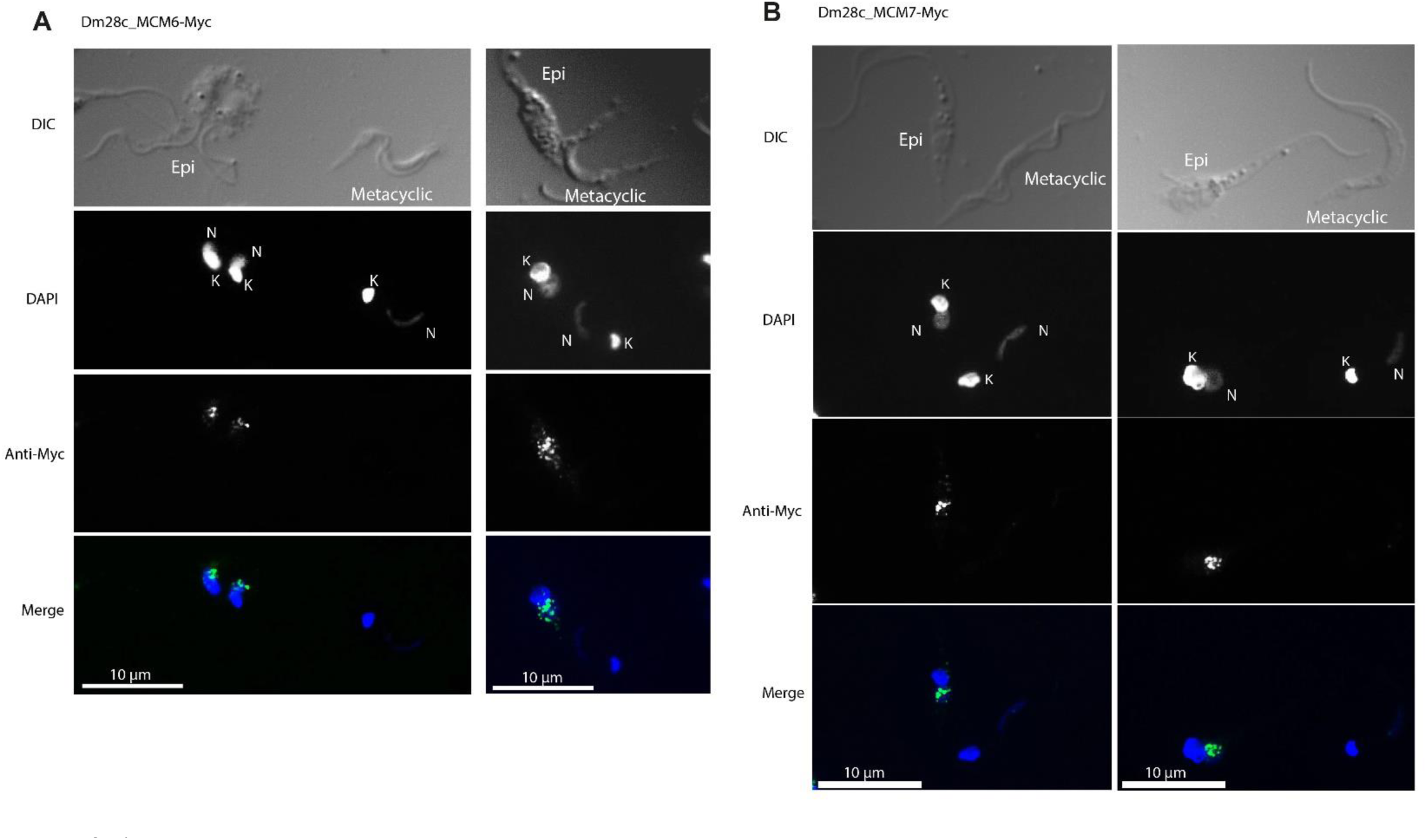
Immunofluorescence imaging of *T. cruzi* Dm28c cells after metacyclogenesis. Epimastigotes from the MCM6-Myc and MCM7-Myc cell lines (Dm28c strain) were subjected to metacyclogenesis. Immunofluorescence staining was performed using an anti-Myc antibody. The images show both differentiated metacyclic trypomastigotes and non-differentiated epimastigotes. (**A**) MCM6-Myc cell line and (**B**) MCM7-Myc cell line. Anti-Myc staining is shown in green, DAPI-stained DNA in blue, and Differential Interference Contrast (DIC) images are included. The white scale bar represents 10 µm. N denotes the nucleus, K the kinetoplast.

